# Anti-cancer Potential of *Moringa oleifera* on BRCA1 Gene: Systems Biology

**DOI:** 10.1101/2020.12.19.392423

**Authors:** Toheeb A. Balogun, Kaosarat D. Buliaminu, Onyeka S. Chukwudozie, Zainab A Tiamiyu

## Abstract

Breast Cancer has always been a global challenge that is prevalent among women. There is a continuous increase in the high number of women mortality rates as a result of breast cancer and affecting countries at all levels of modernization. Women with high-risk factors including hereditary, obesity, and menopause have the possibility of developing breast cancer. With the advent of radiotherapy, chemotherapy, hormone therapy, and surgery in the treatment of breast cancer, there has an increased number of breast cancer survivors. Also, the design and development of drugs targeting therapeutic enzymes are helping to effectively treat the tumor cells at an early stage. However, long term use of anti-cancer drugs has been linked to severe side effects. This research aims to develop potential drug candidates from *Moringa oleifera* which could serve as anti-cancer agents. *In silico* analysis using Schrödinger Molecular Drug Discovery Suite and SWISS ADME was employed to determine the therapeutic potential of phytochemicals from *M. oleifera* against breast cancer via molecular docking, pharmacokinetic parameters, and drug-like properties. The result shows that Rutin, Vicenin-2, and Quercetin-3-O-glucoside have the highest binding energy of −7.522, −6.808, −6.635kcal/mol respectively in the active site of BRCA1. The essential amino acids involved in the protein-ligand interaction following active site analysis are ASN 1678, ASN 1774, GLY 1656, LEU 1657, GLN 1779, LYS 1702, SER 1655, PHE 1662, ARG 1699, GLU 1698, and VAL 1654. Thus, we propose that bioactive compounds from *M. oleifera* may be potential hit drug candidates against breast cancer.

## INTRODUCTION

Breast cancer is the leading cause of death in women around the world. Several factors contribute significantly to the increased risk of breast cancer which includes oral contraceptives, obesity, menopause, and elevation in serum estradiol concentration [1]. Ductal carcinoma is the most common type of breast cancer which developed from the ducts. Cancerous cells developing from lobules are called lobular cells [2]. Breast cancers are mostly diagnosed by physical examination by a healthcare provider or the use of mammography [3]. High occurrence of breast cancer has been reported to be prevalent in white women within the range of forty years and above [4].

Breast cancer gene 1 (BRCA-1) also called caretaker gene is a tumor suppressor gene that functions in cell cycle regulation, DNA repair mechanism, and other metabolic processes [5][6]. The BRCA-1 proteins interact with other essential proteins necessary in DNA replication and repairing of double-stranded DNA breaks [7]. It contains 1863 amino acid residues and helps to inhibit the proliferation of cells lining the milk ducts of the breast. Thus, BRCA-1 does not contribute to the pathogenesis of breast cancer. However, mutations in the sequence of the Breast Cancer gene can consequently increase the risk of breast cancer [8]. Mutations evolved when the genetic makeup of an individual becomes damaged via exposure to environmental factors including ultra-violet light, ionizing radiation, and genotoxic chemicals [9]. When the BRCA-1 is mutated, it cannot efficiently repair the broken DNA, thereby, the prevention of breast cancer will be hampered [10].

There are several treatment methods available for breast cancers but hormone-blocking agents, chemotherapy and monoclonal antibodies are the most commonly used [11] [12]. Hormone receptors (estrogen ER+ and progesterone PR+ receptors) are a therapeutic target in breast cancer. Drugs such as tamoxifen and anastrozole act by blocking the hormone receptors [13]. Several medicinal plants such as *Camptothecan acuminate, Catharanthus roseus, Taxus brevifolia*, and many others have been used as anti-cancer therapy [14].

*Moringa oleifera* which belongs to the family of *Moringaceae* has been reported to possess beneficial pharmacological properties such as anticonvulsant, antimicrobial, anticancer, and antiviral [15]. The extracts (phytochemicals) from the leaves, seeds, bark, and flowers of *M*.*oleifera* have been used in the treatment of several chronic diseases including hypercholesterolemia, high blood pressure, diabetes, insulin resistance, non-alcoholic liver disease, cancer, and inflammation [16]. Bioactive compounds of *M. oleifera* shows inhibitory potential against cancerous cell line by inhibiting proliferation of carcinoma cells and malignant astrocytoma cells [17] [18]. In this study, *in silico* analysis via: molecular docking and pharmacokinetic profiles were employed to screen the library of bioactive compounds from *M*.*oleifera* to determine their anticancer property.

## MATERIALS AND METHOD

### LIGAND PREPARATION

The phytochemicals of *M. oleifera* were retrieved from published literature [15] and their crystal structures were downloaded from the PubChem database (https://pubchem.ncbi.nlm.nih.gov/). The PubChem Compound Identification Numbers (CIDs) for each ligands are Rutin (CID: 5280805), Vicenin-2 (CID: 5280805), Quercetin-3-o-glucoside (CID: 5748594), Chlorogenic acid (CID: 1794427), Gallic acid (CID: 370), Sinalbin (CID: 656568), Isoquercetin (CID: 5280804), Astragalin (CID: 5282102), Quercetin (CID: 5280343), Ferulic acid (CID: 445858), Myricetin (5281672), Kaempferol (CID: 5280863). The ligands were prepared using the LigPrep module of Glide tool by utilizing the OPLS 2005 force field [19].

### PROTEIN PREPARATION

The crystal structures of the BRCA-1 (PDB ID: 4OFB) was retrieved from Protein Data Bank (https://www.rcsb.org/) in complex with co-crystallized ligands. The protein was prepared using ProteinPrep Wizard of Maestro interface (11.5) by adding missing hydrogen atoms. Furthermore, the metal ionization was corrected to ensure formal charge and force field treatment. The protein was optimized and refined for docking analysis [20] [21].

### MOLECULAR DOCKING

The docking analysis was conducted using the Glide tool from Schrodinger molecular drug discovery suite (version 2017-1). The grid was generated using the receptor grid generation module of the Glide tool. The coordinate (x, y, z) of the grid was centered to −9.07, 27.02, and - 0.91 respectively. The refined *M*.*oleifera* ligands were docked into the active site of BRCA-1. The energy calculation was achieved using the scoring function of the Glide tool. The drug-like properties of the compounds were evaluated using the QikProp module and SWISS ADME Web tool following Lipinski’s rule of five [22].

## RESULTS AND DISCUSSION

Molecular docking was employed to carry out the virtual screening of the library of phytochemicals from *Moringa oleifera* against the targeted protein (BRCA-1). The phyto-compounds of *M*.*oleifera* were ranked according to their binding poses and energy calculations. The compounds were further subjected to pharmacokinetic study to predict their drug-able properties. The molecular docking analysis which includes: binding affinity (Kcal/mol) predication, the interaction of the ligands within the binding pocket of BRCA-1, and their pharmacokinetic study was shown **(Table 1)**. Each ligand was analyzed using Lipinski’s rule of five (ROF), the result confirms that the ligands ROF with few violations. The ligand docking shows how the phyto-compounds bind effectively with BRCA-1. Visualization of the protein-ligand complex was carried out using the surface module of the Glide tool (Figure 1). The interaction between the compounds and BRCA-1 identified the amino acid residues involved in the interaction as well as the position of each amino acid residues in their ligand-binding site. The interaction was associated with a structure-based drug design depicting protein-ligand interaction.

**Table 1:**
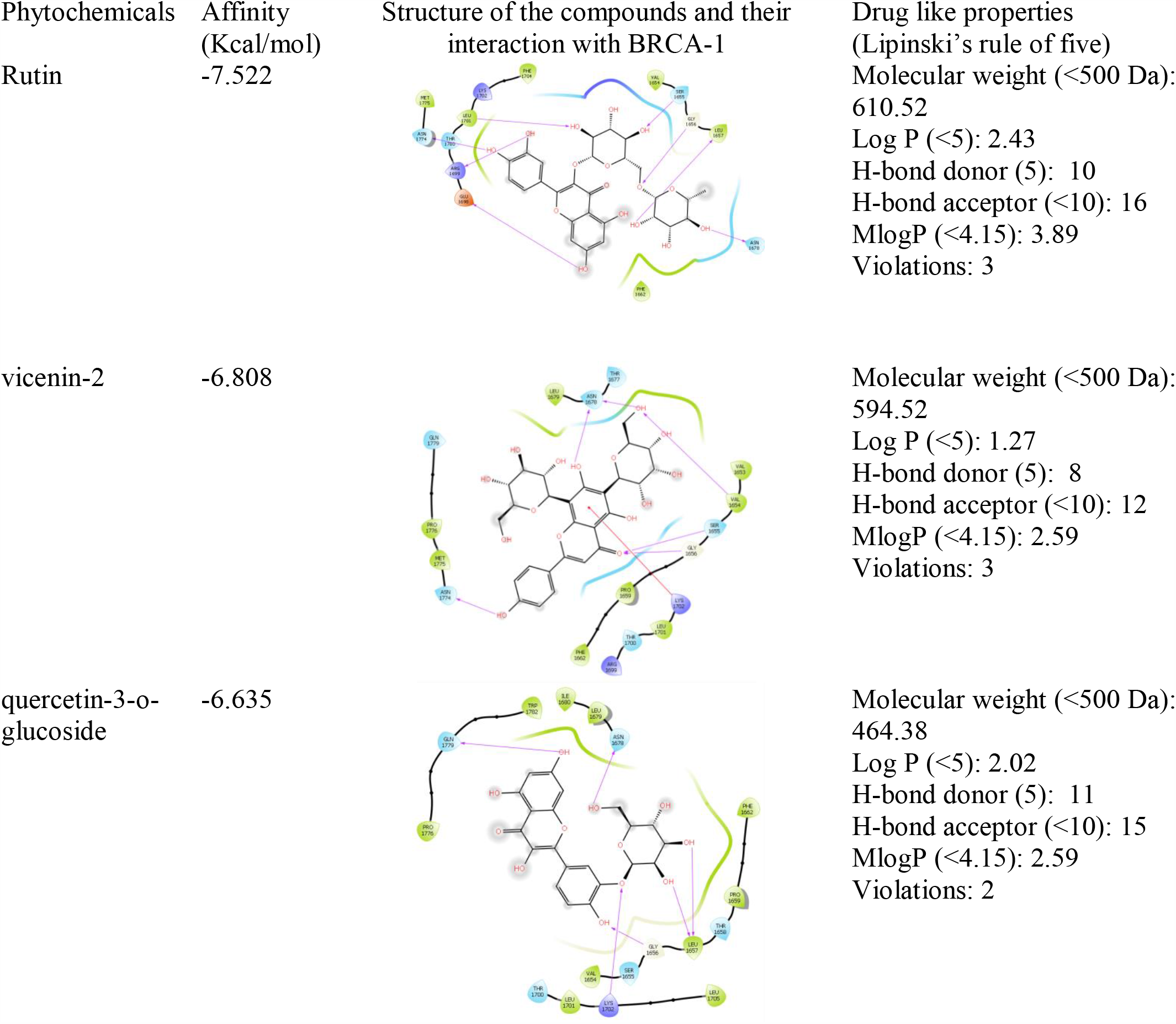
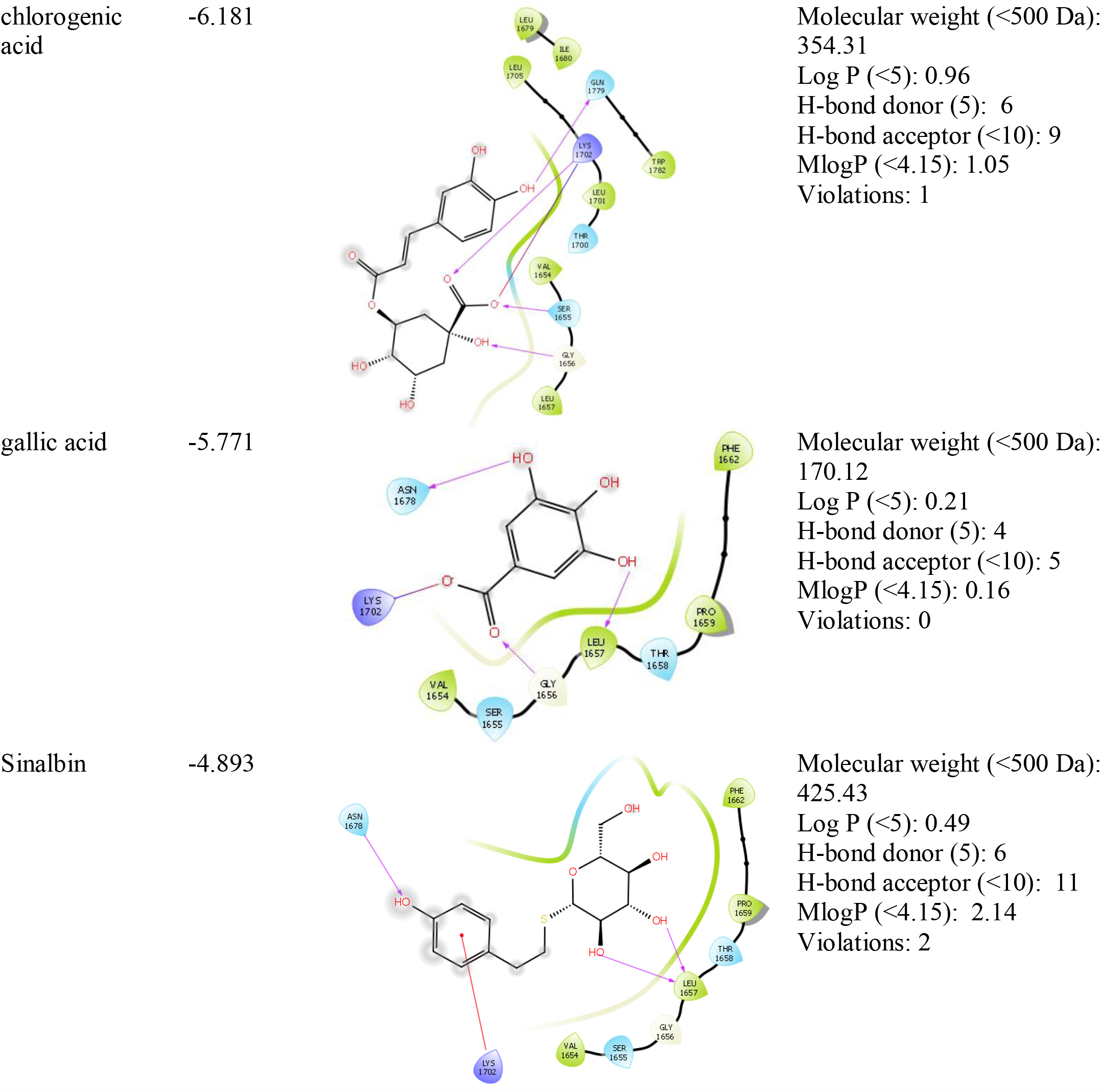
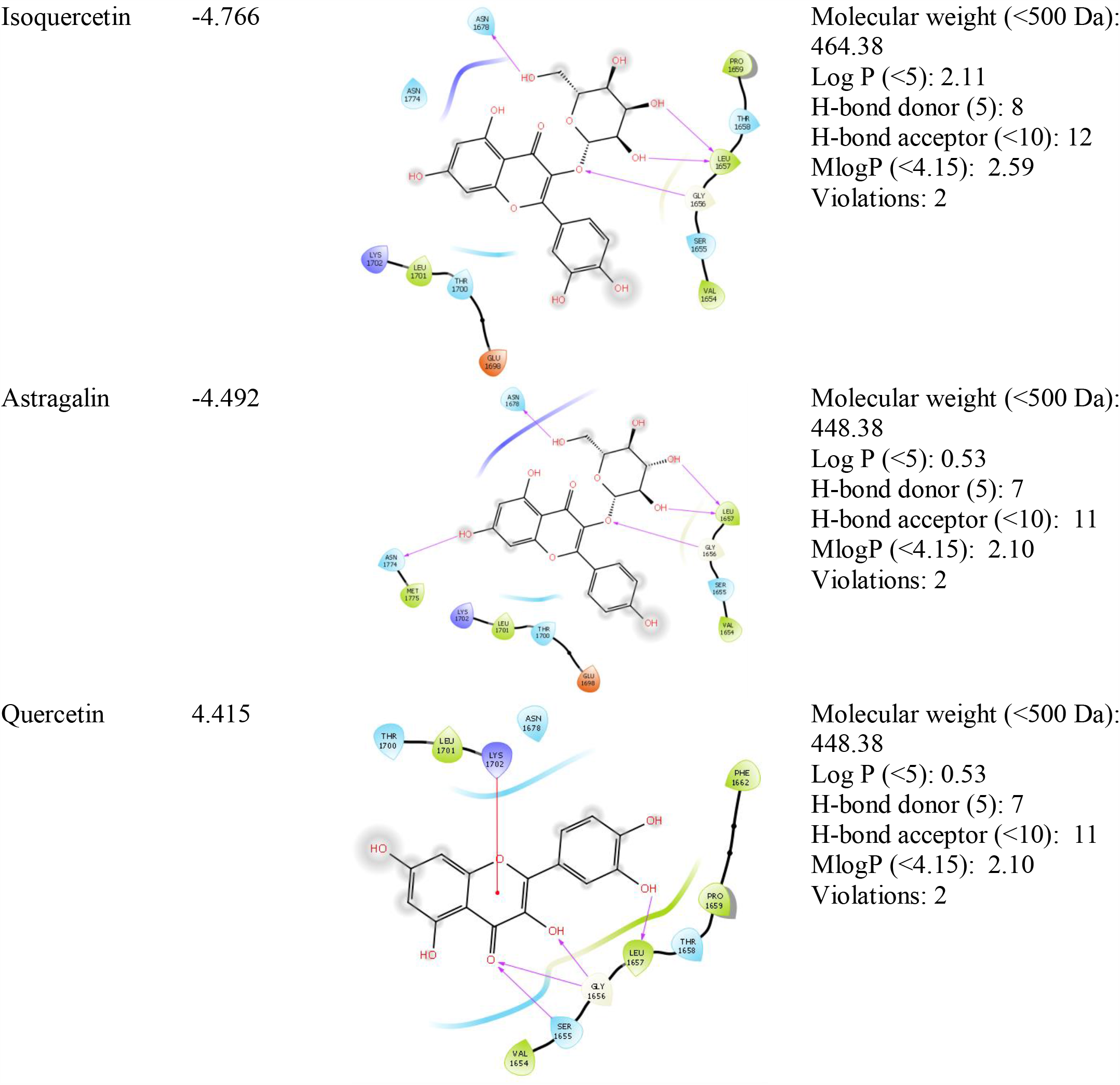
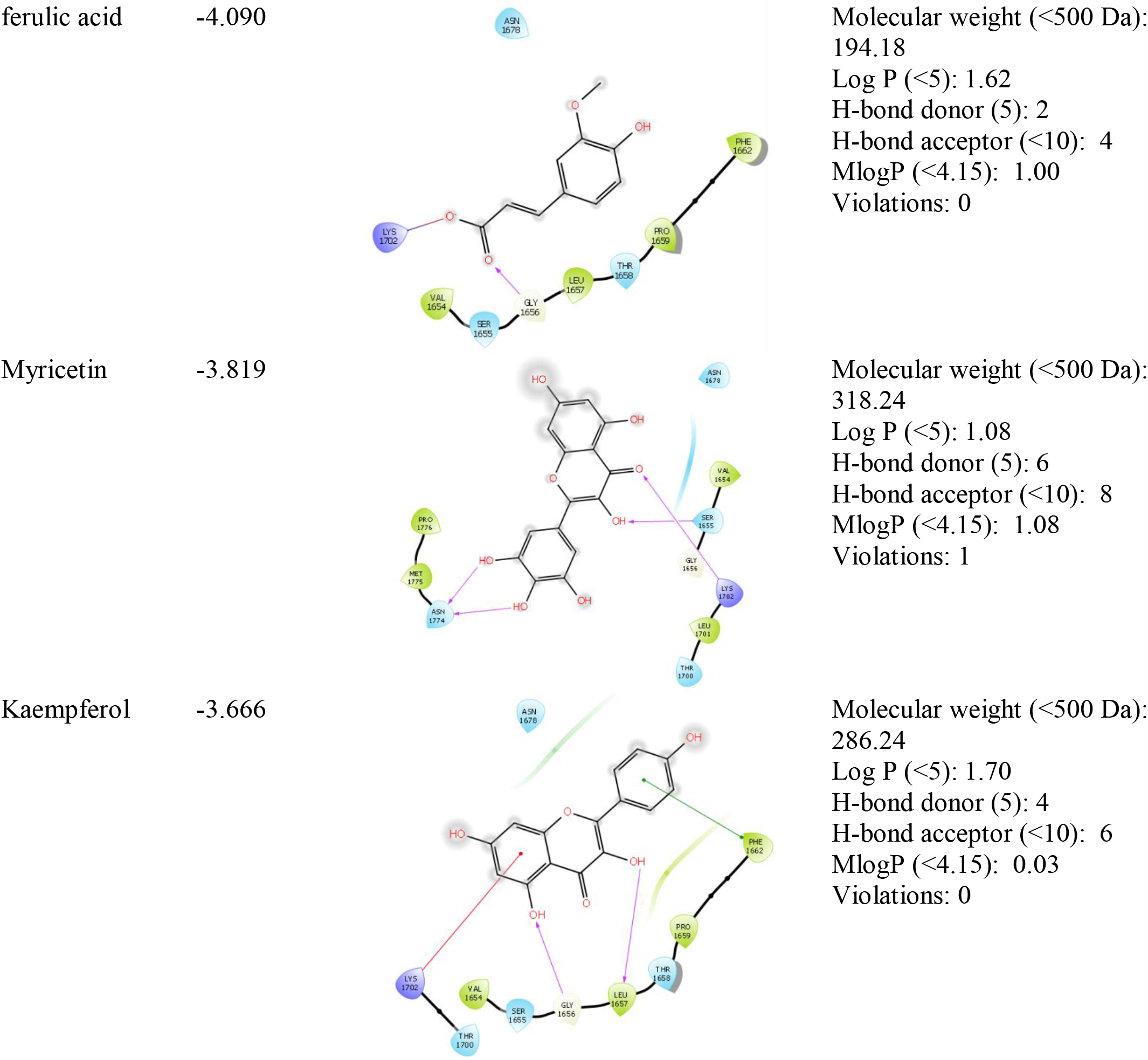
Docking results of phytochemicals from *M. oleifera* in terms of binding affinity (kcal/mol), the interaction of the compounds with BRCA-1, and the drug-like properties.

**Figure 1:**
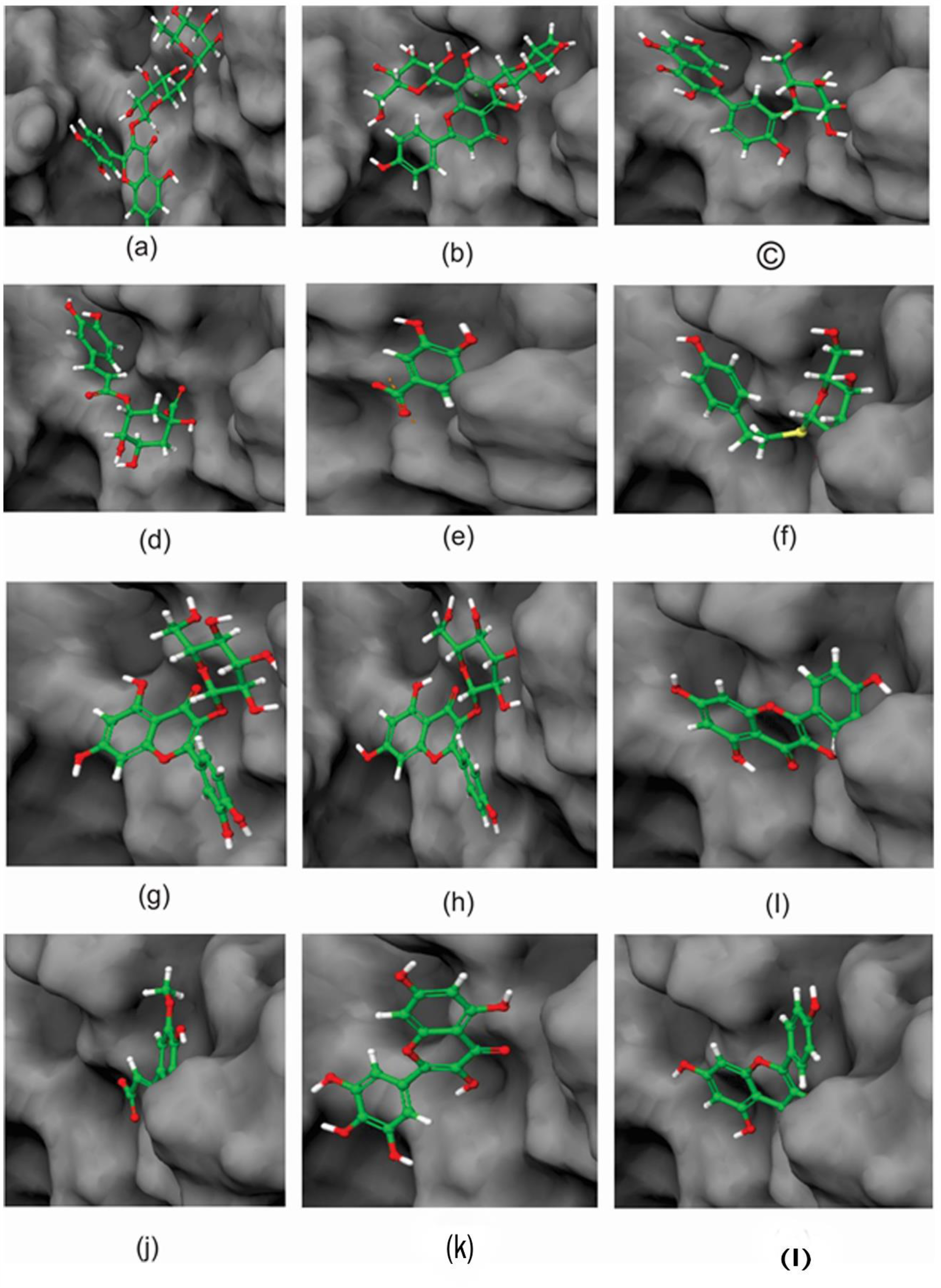
Visualization of docking results showing binding of **(a)** Rutin (**b)** vicenin-2 **(c)** quercetin-3-o-glucoside **(d)** chlorogenic acid **(e)** gallic acid **(f)** sinalbin **(g)** Isoquercetin **(h)** Astragalin **(i)** quercetin **(j)** ferulic acid (**k)** Myricetin (**l)** kaempferol with BRCA-1.

The molecular docking demonstrates interactions such as hydrophobic, pi-pi stacking, hydrogen bonding, and many others between the protein and the ligands. BRCA-1 was co-crystallized with a natural inhibitor which defines its active site. This allows the binding of the ligands into the protein-binding domain. The primary amino acids involved in the protein-ligand interaction following active site analysis are ASN 1678, ASN 1774, GLY 1656, LEU 1657, GLN 1779, LYS 1702, SER 1655, PHE 1662, ARG 1699, GLU 1698, and VAL 1654. The phytoconstituents show a favorable interaction with BRCA-1. Rutin (a flavonoid), following extra precision docking against BRCA-1 shows hydrogen bonding interaction and pi-pi stalking with amino acid residues LEU 1701, ASN 1774, ARG 1699, GLU 1698, ASN 1678, LEU 1657, SER 1655 and a binding affinity of −11.769Kcal/mol. The toxicity study of Rutin confirms that it has a low bioavailability, binds firmly to the human serum albumin, high metabolic rate, and can be easily excreted. Vicenin-2 exhibited promising ligand interaction when complexed with BRCA-1. It binds with an energy of -6.808kcal/mol by hydrophobic interaction with VAL 1654. The docking of Quercetin-3-o-glucoside with BRCA-1 exhibited a glide score of −6.635Kcal/mol by forming five hydrogen bonds with ASN 1774, GLY 1779, ASN 1678, GLY 1656, and Ser 1655 accompanied with pi-pi stacking at amino acid residue LYS 1702. The compound, Chlorogenic acid binds well with the targeted protein with an affinity of −6.181Kcal/mol. There was a favorable interaction of gallic acid, sinalbin, and Isoquercetin against BRCA-1 with a binding energy of −5.771, −4.893, and 4.766 respectively. The drug-like properties of gallic acid demonstrated that it does not violate Lipinski’s rule of five with a promising therapeutic potential. Isoquercetin interacts with an amino acid at GLY 1656. The pharmacokinetic profiles of Astragalin and Quercetin adhere to the ROF with only two violations and docking scores of 4.415, −4.090Kcal/mol respectively. Ferulic acid and Myricetin have a binding energy of −4.090 and −3.819Kcal/mol respectively when complexed with the targeted protein. Kaempferol has drug-like properties without violating Lipinski’s ROF and binding energy of −3.666Kcal/mol. The various interactions between the ligands and BRCA-1 that, the compounds may be potential anticancer agents.

## CONCLUSION

Several anti-cancer drugs such as tamoxifen, anastrozole, exemestane, have been developed and are effective but posed serious side effects following long-time use including liver toxicity, cardiovascular diseases, and many others. In this study, we utilized computational modeling techniques to predict the inhibitory potential of *M. oleifera* against BRCA-1. The binding of the compounds with BRCA-1, toxicity, and drug-like property as confirmed by docking analysis shows that the *M*.*oleifera* ligands are promising anticancer agents. Following the screening of the Phyto-compounds from *M*.*oleifera* by docking technique, Rutin, was found to exhibit the highest degree of interaction and binding affinity with BRCA-1 accompanied by favorable drug-like properties. Thus, we proposed that the phytochemicals from *M. oleifera* may be potential BRCA-1 inhibitors. Further biochemical analysis such as *invitro and invivo* study is required to establish the pharmacological properties of the compounds.

## Notes

### Competing Interest Statement

The authors have declared no competing interest.

